# Learning a mechanical growth model of flower morphogenesis

**DOI:** 10.1101/2025.02.27.640522

**Authors:** Argyris Zardilis, Alexandra Budnikova, Henrik Jönsson

## Abstract

Morphogenesis, the process by which organs take their shape, is a fundamental problem in biology and requires a complex combination of genetic, biochemical, and mechanical signals. The data that we collect for this process are usually multi-scale and multi-modal, typically image timeseries of tissue morphology and transcription data. As the data becomes larger and more diverse, the harder it becomes to gain meaningful understanding in terms of theoretical models. Here, and tackling this problem in the developing flower organ, we learn a mechanical model of growth starting from combined morphology and transcription data across the tissue by firstly learning and connecting independent representations for the two spaces and secondly refining the representation using the mechanical state of the tissue to learn a mechanical growth model. This approach can link multiple scales, in this case proposing the physical action of genes, and can act as a vehicle to accommodate diverse data and hypotheses in developmental biology of plants and beyond.

## 1 Introduction

Morphogenesis, the process by which organs and organisms acquire their shape, requires the integration of genetic, biochemical and mechanical signals. The data that we collect for this process is therefore multimodal and multi-scale, typically image time series of the morphology of the developing tissue and some measure of the (spatial) expression of genetic and hormonal components. As the data grows in size and diversity, integrated analysis of these datasets and incorporation into mechanical theories of growth becomes extremely challenging to do manually Berens et al. (2023). We address this problem in sepals – the first organs of the flower, emerging in the outer whorl of the flower bud and transitioning from outgrowths to folded structures – as an example system of complex organogenesis Roeder (2021). In previous work we have developed an ‘atlas’ of spatial gene expression for this process at cellular resolution, integrating a number of genetic components in the same reference time series Refahi et al. (2021) (Figure 1a). Here we use this dataset to infer a mechanical model of growth to propose possible physical mechanisms of action of molecular species by (i) computing separate representations for the morphology and genetic views of the dataset and learning a mapping between them to identify genetic components more likely to have contributed to morphological characteristics of the tissue and (ii) inferring a linear mechanical model of growth for this process using the identified genetic components from (i), calculated cell growth rates and computed mechanical state. This representation integrates genetic and mechanical signals to give a more comprehensive picture of the process.

**Figure 1:**
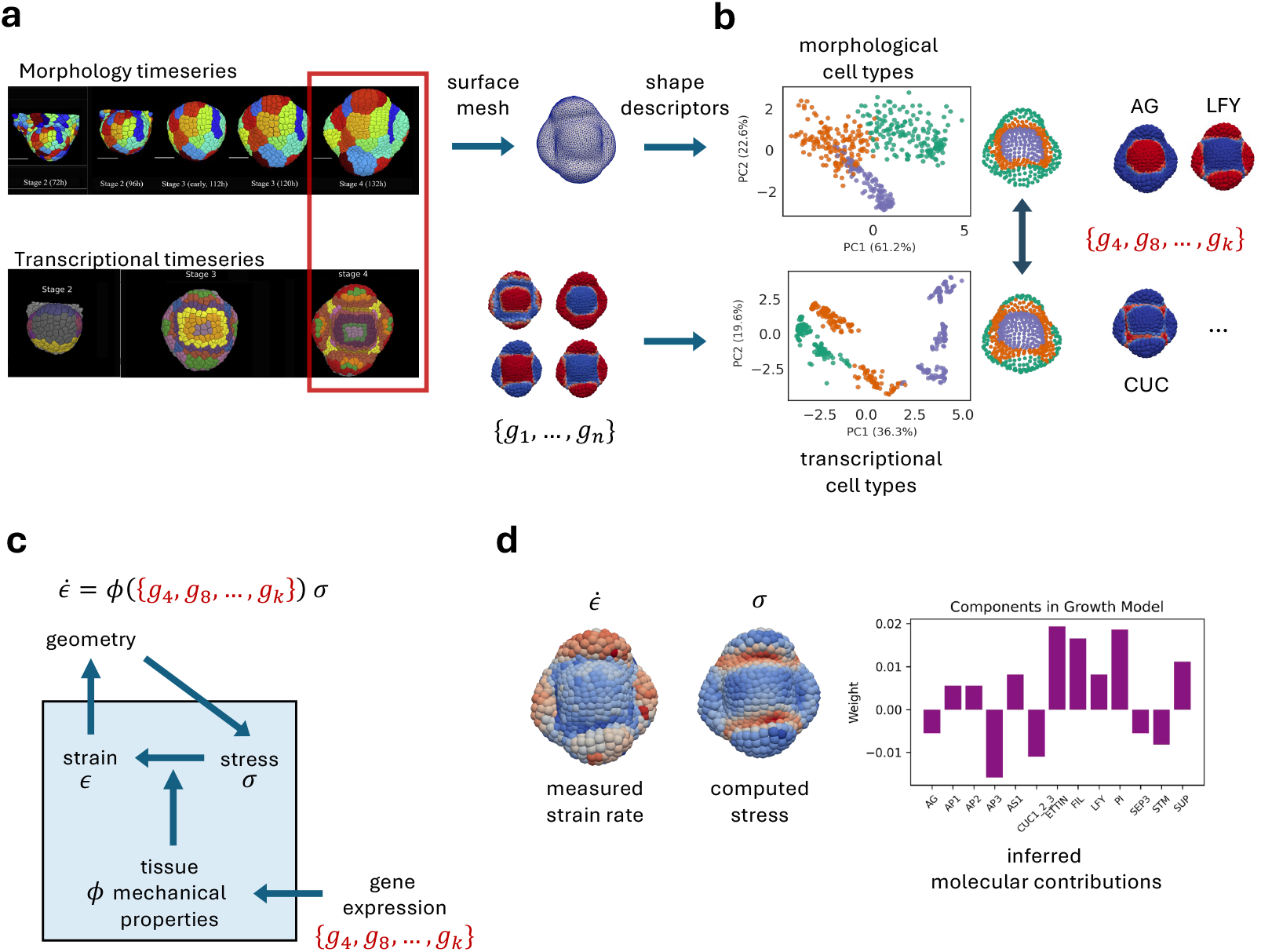
a. Morphological characteristics are computed from the last time point of an imaging time series of the developing organ and are b. mapped to spatial gene expression to get the genes most correlated to morphology c. Schematic of the mechanical growth model including the proposed influence of genetic factors d. Inferred molecular contributions to growth using the computed strain rates and mechanical stress of the tissue. Images of the time series in a. are reproduced from Refahi et al. (2021).

## 2 Results

### 2.1 Data representation

One way to infer the causality of genes in (flower) development is through perturbations, so if we perturb a gene and see an effect on the organ identity or morphology of the tissue, we can link them causally. This led to the identification of important developmental genes and models of their action in flower morphogenesis (ABC model; Bowman & Moyroud (2024); Coen & Meyerowitz (1991); Bowman et al. (1989)). Since plant cells are non-migrating, their action should be local so here we will use spatial correlation between morphological characteristics and gene expression as a proxy of causality. From the reference time series we focus on the last time point as it is the first stage where we can see distinct morphological aspects of the sepals (Figure 1a).

To compute morphological characteristics we first extract a surface mesh from the 3D image stack and then for each point on the mesh compute the principal curvatures at that point, *κ*_1_ and *κ*_2_, and other curvature descriptors based on those; gaussian curvature (*κ*_1_*κ*_2_), mean curvature ((*κ*_1_+*κ*_2_)*/*2), and deviatoric curvature ((*κ*_1_ *− κ*_2_)*/*2). These curvature metrics are transferred to the tissue cells and combined with cell volume. With the chosen measures we tried to capture the local geometry around a cell but there are more morphological features that could potentially be interesting like the shape of the cell itself and other more series-based descriptions of shape (based e.g. on spherical harmonics as in Pönisch et al. (2024)). On the other hand the spatial gene expressions are defined in the atlas per cell and are spread using a diffusion process to make them continuous as they were initially binary. Using PCA for dimensionality reduction and a Gaussian Mixture model for clustering, these features seem to capture the main regions and cell types at this stage, the central stem cell region, the developing organs at the four poles, and the boundary between them (Figure 1b). We next use logistic regression to learn functions between the morphological characteristics and each of the 25 genes across the cells in the dataset. From the 25 genes, 13 were well predicted from morphology (balanced accuracy score *>* 0.75) including some well known markers for cell states, like CUC (boundary), LFY (sepals) and AG (central region) (Figure 1b, Alvarez-Buylla et al. (2010)).

### 2.2 Model inference

We have identified a statistical relation between morphology and gene expression but how could the genes have caused the morphology? Since plant cells are surrounded by a cell wall, morphogenesis becomes purely a problem of growth and the role of cell wall mechanics is especially apparent. Assuming a linear mechanical model, the irreversible growth (strain) rate of the tissue at any point is given by: 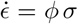 where *σ* is a measure of the mechanical stress and *ϕ* is a measure of the material properties and addition of new wall material (extensibility) Lockhart (1965); Ortega (1985). We can think of growth as the balance between the amount of force and the amount of resistance by the tissue. Here, for simplicity, we omit the ‘yield’ stress by which growth only occurs after a threshold 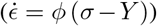. One proposed way that genes affect this process locally is by changing the material properties of the tissue to allow either more or less resistance to the mechanical forces that then gives rises to differential growth, 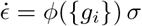 (Figure 1c).

Since we have live imaging and tracking of the cells, we can compute their volumetric growth rate 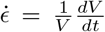, computed as in Refahi et al. (2021)) and we can also computationally infer the mechanical state of the cells (*σ*, computed as in Bozorg et al. (2014)). The inference problem can then be posed as:

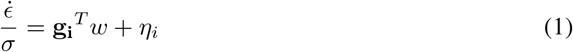

where we assume a linear effect of the genes on the mechanical properties. The weights to learn *w* can then be interpreted as the contributions of each gene to the mechanical properties and consequently growth via mechanical stress (Figure 1d). Practically, in solving 1 we also used the *l*^2^-norm of the weights to balance the fitting with model simplicity:

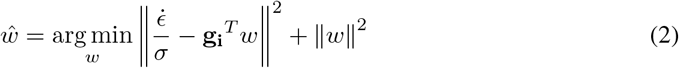

## 3 Discussion

We have proposed a method to infer a mechanical growth model to propose physical action of genes, connecting molecular, mechanical and morphology data to get a more comprehensive picture of development in a complex organ. As with any model there are simplifications. Plant growth is unlikely to be linear and volumetric growth, like we have assumed, misses important aspects of growth like direction and anisotropy. There have been many other components not included here that have been proposed to influence aspect of growth like the microtubule networks in cells and the hydrology of the tissue Oliveri & Cheddadi (2025); Hamant et al. (2008). Growth can be completely captured tensorially, which can capture all its aspects, but then the biggest problem might be computing these quantities from data. Nevertheless, we still argue that our methods despite its simplifying assumptions gave us a way to integrate signals in this process and a way to propose a physical mechanism of action.

As for our data, we took advantage of the integrated dataset but this might not be the case in general. It is more likely that data comes in as, for example, single-cell sequencing data. This makes the integration harder but it should be possible to develop similar methods for such data. Moreover we ignored previous time steps in our time series data but computing trajectories in the two spaces and linking those will be valuable and come closer to causality. We can imagine, for example, even learning the terms of the physical theoretical model to further explore the space of hypotheses Kamienny et al. (2022); Maddu et al. (2021).

In conclusion, our proposed method is a powerful way to integrate data from multiple processes in development. These methods can be developed further to act as a vehicle for integrating diverse hypotheses going beyond genes in developmental biology and should carry to other developmental processes in plants and beyond Özpolat et al. (2025).

